# Introduction of Dicistrovirus IRESs into UAS/SV40-polyA constructs results in premature polyadenylation and strong overexpression of the upstream ORF in *Drosophila* animals

**DOI:** 10.1101/2023.10.04.560905

**Authors:** Peter V. Lidsky, Sergey E. Dmitriev, Raul Andino

**Affiliations:** Department of Microbiology and Immunology, University of California San Francisco, San Francisco, 94158, CA; Belozersky Institute of Physico-Chemical Biology, Lomonosov Moscow State University, Moscow 119234, Russia

## Abstract

To evaluate the properties of insect virus internal ribosomal entry sites (IRESs) for protein expression in *Drosophila*, we have introduced Cricket Paralysis virus (CrPV) and Drosophila C virus (DCV) IRESs into UAS/SV40-polyA vector. We found that introduction of IRESs induce premature polyadenylation, resulting in both truncation of the mRNA, and an increase in mRNA levels of approximately 40-fold. The increase in mRNA levels was accompanied by increased resistance to nonsense-mediated mRNA decay (NMD)-mediated degradation. Our results suggest that premature polyadenylation increases mRNA stability in the SV40 polyadenylation site-containing constructs, suggesting a novel method for robust overexpression of transgenes in *Drosophila*.

## Introduction

*Drosophila melanogaster* is an important model organism in scientific and biotechnological studies. The regulation of transgene expression is critical in many scientific and biotechnological endeavors involving *Drosophila*, as demonstrated in various studies (Jennings, 2011; Mirzoyan et al., 2019; Tolwinski, 2017). The ability to regulate the expression of transgenic proteins is essential to increase signals intensities when using fluorescent reporters (Bothma et al., 2018; Pfeiffer et al., 2012, 2010), to perform overexpression experiments (Schittenhelm et al., 2010), and for other applications.

Insect cells are widely used for protein production due to their ability to produce high yields of recombinant proteins (Ikonomou et al., 2003; Maiorella et al., 1988; Palomares et al., 2004). Genetically modified insects are also used in industry to produce nutritional additives or bioactive proteins, and scaling up transgene expression may be useful in such contexts (Patel et al., 2019; van Huis and Oonincx, 2017; Wang and Shelomi, 2017).

Virus-derived Internal Ribosomal Entry Sites (IRESs) have been commonly employed in mammalian systems for protein expression (Elroy-Stein et al., 1989; Jaafar and Kieft, 2019; Jan, 2006; Mailliot and Martin, 2018; Sorokin et al., 2021; Terenin et al., 2017). Interestingly, though IRESs are present in many insect viruses and are included as part of their protein synthesis process, they have not yet been effectively utilized for insect protein expression (Sasaki and Nakashima, 1999; Wilson et al., 2000). In this study, we have placed two IRESs (Jan, 2006) derived from insect viruses – one from Cricket Paralysis virus (CrPV) and the other from Drosophila C virus (DCV) into bicistronic constructs and produced transgenic flies encoding these constructs. Both IRESs were unable to induce detectable translation of the second cistron in healthy tissues of fruit flies. Surprisingly, these IRES-encoding constructs generate shortened polyadenylated transcripts likely due to early transcription termination induced by the presence of cryptic polyadenylation signals within the IRES sequences. Truncated transcripts were present at extremely high levels, resulting in a very strong expression of proteins encoded by the ORF upstream of the IRES and retained in these transcripts. Our results indicate that while the full-length constructs are subject to degradation by nonsense-mediated mRNA decay (NMD), short transcripts are not, explaining the overexpression effect. Furthermore, our results suggest that the IRES of CrPV may contain an element that protect the full-length transcript from NMD-mediated degradation.

## Results

### Sequences within the CrPV and DCV IRESs enable overexpression of foreign proteins encoded upstream of these IRESs

To analyze the IRES activities in *Drosophila* organisms, we engineered bicistronic constructs with CrPV and DCV intergenic IRESs. In these bicistronic constructs, the first open reading frame, encoding red fluorescent protein mCherry, is translated in a cap-dependent manner, while the second cistron, encoding EGFP, is controlled by either the CrPV, or DCV IRES. As a control we used a monocistronic expression cassette encoding mCherry-EGFP fusion protein, expressed by a cap-dependent mechanism. All these constructs were transcribed from an UAS promoter (Brand and Perrimon, 1993), and the mRNA polyadenylation was mediated by the SV40 polyadenylation signal at the 3’ end of expression cassettes (**Fig. 1A**). All constructs were inserted into the 86Fb-attP attachment site on chromosome 3 using ΦC31 integrase (Bischof et al., 2007). Transgenic flies were crossed with the *da:Gal4* driver line to test IRES-mediated expression. In the majority of fly tissues, EGFP expression driven by IRES was not detectable (**Fig. 1B**, middle row). IRES-dependent translation was observed under conditions mimicking the integrated stress response. If constructs were co-expressed with constitutively active mutant of GCN2 kinase (Bjordal et al., 2014) they were able to produce detectable EGFP signal (**Fig. 1D**), as it was described previously for the IRES from Black Queen Cell virus (Lidsky et al., 2023). Therefore, CrPV and DCV IRESs, when placed in bicistronic constructs, were capable to initiate translation. In IRES-containing constructs, we found that mCherry fluorescence levels were greatly elevated compared to the monocistronic control (**Fig. 1B**, upper row). The red color of the flies was so intense that it could be seen without any equipment in natural daylight - a phenomenon made possible by the high levels of overexpression (**Fig. 1B**, bottom row). Northern analysis revealed the presence of an unexpected shorter form of mCherry-encoding transcript that expressed at apparently high levels in IRES-containing constructs (**Fig. 1C**).

**Fig. 1.**
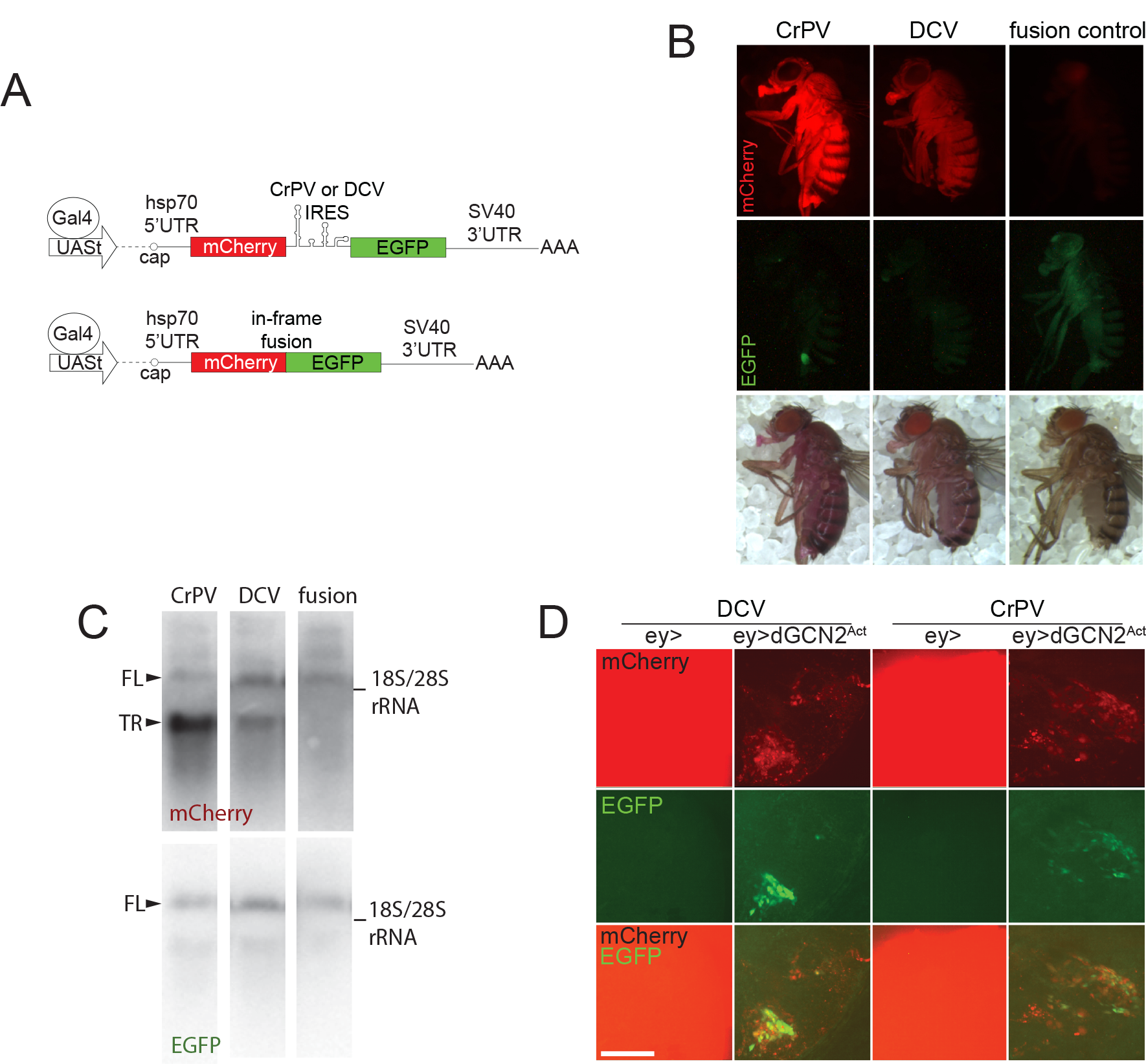
Overexpression of the first cistron in IRES-containing constructs. **A**. Scheme of the constructs used in this study. **B**. Flies homozygous by *da:Gal4* and a corresponding construct were examined with epifluorescent microscopy. All images are taken under the same conditions and with the same exposure. Images in the bottom row are taken in the daylight with no excitation light. No green fluorescence was detected in animals harboring IRES-encoding constructs. The green dot in the CrPV sample is a piece of food attached to the fly body. **C**. Northern blot analysis of total RNA fraction isolated from flies expressing *da:Gal4* and indicated constructs in single copies. FL denotes the full-length transcript, TR denotes the truncated transcript. rRNA was used as a size marker. **D**. IRES-containing constructs can produce EGFP upon induction of integrated stress response. We co-expressed bicistronic constructs together with constitutively active mutant of GCN2^F751L^ kinase in the optic lobes of the larval brains. 3^rd^ instar larvae were collected for this panel. GCN2 ^F751L^ expression drastically reduces cell viability, the expression of mCherry and induced expression of EGFP. Images are made with the same exposure. Bar is 100 μm.

### The transcripts produced with IRES-containing constructs are less susceptible to nonsense mediated decay

The observations described above suggest a model in which overexpression of the mCherry first cistron depends on an increase of truncated mRNA abundance. The ∼700 bp SV40-derived fragment containing polyadenylation cis-acting element is widely used in most UAS-based constructs(Brand and Perrimon, 1993). The lengthy 3’UTR produced with this element makes the transcripts sensitive to NMD (Metzstein and Krasnow, 2006). NMD is a powerful mechanism that prevents the translation of aberrant mRNA transcripts containing premature termination codons (PTCs). The distance between the PTC and the polyA is an important cis-acting signal that triggers NMD in mammals (Behm-Ansmant and Izaurralde, 2006) and in Drosophila (Gatfield et al., 2003). Natural stop codons are typically located within the terminal exon, close to the polyA, while destabilizing nonsense codons lie more upstream. Mutations that affect NMD in *Drosophila*-so-called “photoshop” mutants - have been shown to strongly increase the levels of mRNA derived from constructs harboring SV40 polyadenylation site (Metzstein and Krasnow, 2006). The authors suggest that the NMD machinery recognizes SV40-derived long 3’ UTRs, causing transcript destabilization(Metzstein and Krasnow, 2006; Nelson et al., 2016).

We, thus, hypothesized that NMD reduces expression of our mCherry-EGFP fusion construct (**Fig. 1A**), but this is prevented by the insertion of CrPV and DCV IRESs. To examine this possibility, we crossed NMD-deficient *Upf2*-mutant flies with the animals harboring *da:Gal4* and bicistronic constructs expressing mCherry and EGFP (**Fig. 2A**). The *Upf2*^*25G*^ is a ‘photoshop’ mutation that disrupts normal NMD function, and is associated with late larval or pupal lethality (Metzstein and Krasnow, 2006). However, third instar larvae surviving to this stage could be analyzed for RNA and protein expression (**Fig. 2B-E**). Firstly, we evaluated mCherry levels using a fluorescent microscope (**Fig. 2B**). The mCherry-EGFP fusion construct showed a visible upregulation in expression in the NMD-null background (**Fig. 2B**, compare fusion WT and mut). Conversely, IRES-encoding constructs showed higher expression levels, but a smaller difference between NMD-null and NMD-wt samples (**Fig. 2B**, CrPV, DCV). Next, we measured mRNA levels using RT-qPCR, with two oligo pair-sets that specifically amplify parts of the mCherry (red bars) or EGFP (green bars) coding regions (**Fig. 2C**). The mCherry RNA amounts were significantly higher in the case of the IRES-containing constructs compared to the fusion construct (**Fig. 2C**). For the CrPV-containing IRES construct, these levels were over 40-fold higher than in fusion control. In samples from flies with IRES-containing cassettes, but not in fusion control samples, relative EGFP RNA levels were relatively lower than those corresponding to mCherry (**Fig. 2C**). Importantly, while the expression of the fusion control construct was upregulated in the NMD-null background (**Fig. 2C**, compare fusion WT with mut), the CrPV construct showed very small upregulation detected only with EGFP oligo pair. These findings indicate that the mRNA isoforms containing the second cistron are present at a lower abundance than the first cistron-containing isoforms. To examine whether the IRES sequence results in RNA truncation, we studied the integrity of the bicistronic constructs by Northern blotting as in **Fig. 1C**. Consistently with the RT-qPCR results, IRES samples showed two bands corresponding to mCherry sequences, whereas fusion control samples did not (**Fig. 2D, E**). High molecular weight product corresponded to the bicistronic full-length transcript revealed with both mCherry and EGFP probes, and the low molecular weight one to an RNA containing the first cistron (mCherry) only. The shorter RNA isoforms displayed low sensitivity to NMD inhibition. The full-length product revealed with EGFP probes was upregulated by *Upf2* inactivation in the case of the control fusion construct, but not the CrPV IRES-containing cassette. Full-length transcript developed with mCherry probe also demonstrated visible upregulation but the quantification was not statistically significant, possibly due to high background levels in the mCherry staining. The DCV IRES-containing construct showed both truncated and the full-length transcripts, increased in the *Upf2*-mutant background, while the difference was statistically significant only for full-length isoform. This data suggests that the IRES sequences in bicistronic constructs lead to truncation of a proportion of transcripts produced and also provide a degree of resistance against NMD. Notably, this was more evident for CrPV IRES. To corroborate these results with different genetic tools, were combined eye-specific Gal4 driver with IRES-encoding constructs and RNAi raised against NMD components *Upf1* and *Upf2* (**Fig. 2F, G**, see also **Supplementary Fig S1** for RNAi validation and eye phenotypes). Transgenic flies were used for RNA isolation and qPCR analysis. The obtained results were consistent with previous observations. When NMD was downregulated, the fusion control RNA was increased similarly with both mCherry and EGFP probes. Levels of mCherry RNA in the construct encoding CrPV IRES showed no dependence on NMD activity. Similarly to the results obtained with the *Upf2*^*25G*^ mutation (**Fig 2C)**, the levels of EGFP RNA were increased in samples with suppressed NMD, but this increase was smaller than in fusion control construct (2.4-2.6 fold, compared to 10.5-17.7 fold). Consistently with Fig 2C, DCV IRES caused increase in mCherry and EGFP RNA levels in the absence of NMD. NMD downregulation by RNAi induced a statistically significant increase in both mCherry and EGFP RNA levels. We assumed that RNAi experiment corroborates the phenomena observed in **Fig B-E** indicating NMD as a key player in IRES-mediated stabilization. RNAi combined with CrPV IRES-encoding construct induced a bigger (3.9-4.6 fold) increase in EGFP levels if compared to *Upf2*-mutant (1.3 fold). This difference in effect sizes between two experiments might be attributed to the variability between tissues or developmental stages.

**Fig. 2.**
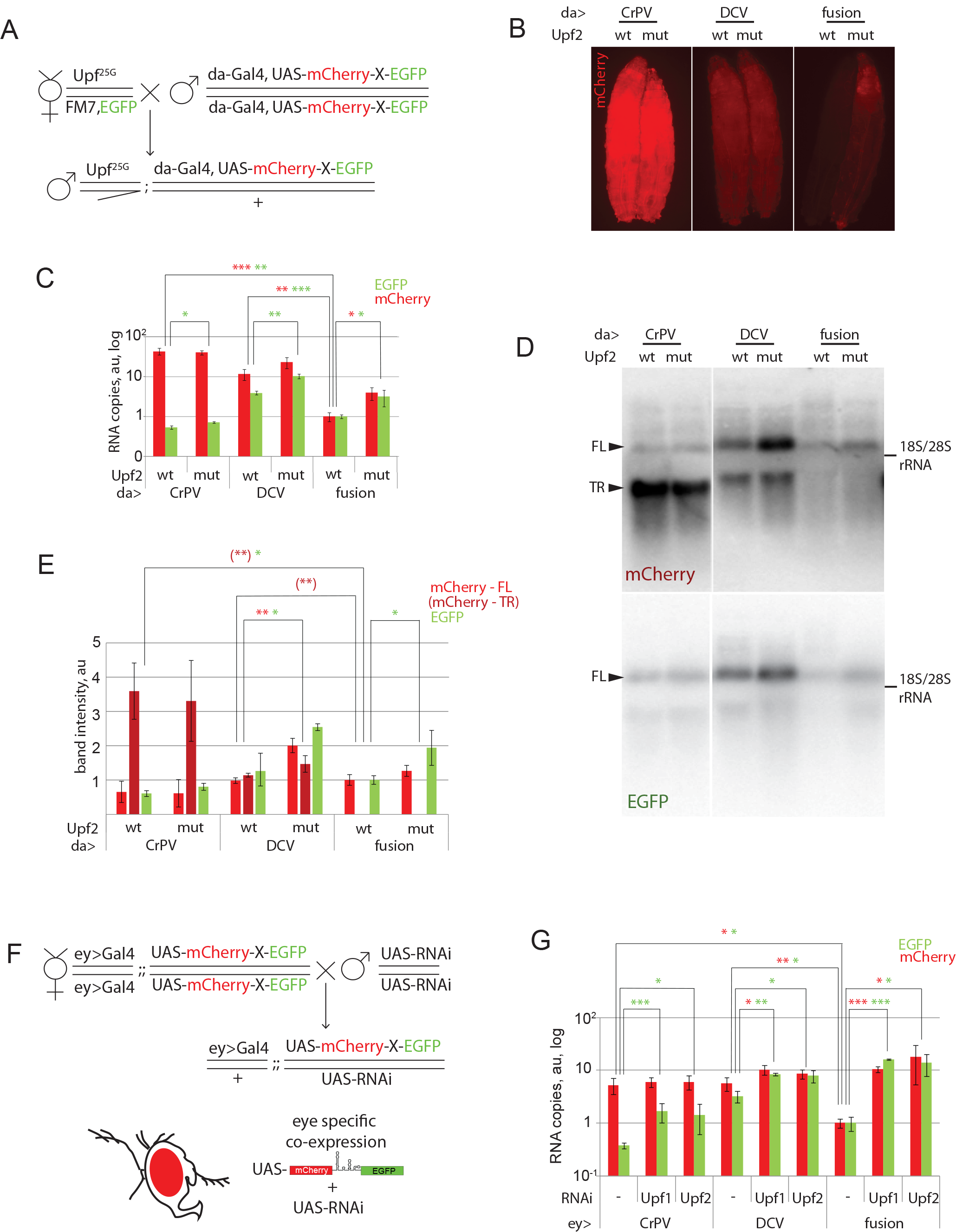
mRNAs produced from IRES-containing constructs display resistance to nonsense-mediated decay. **A.-E.** *Upf*^*25G*^ mutation only slightly affects expression of IRES-containing constructs. **A**. The crossing scheme. Hemizygous males with the *Upf*^*25G*^ mutation or NMD-competent (*yw)* males were analyzed. **B**. NMD-null animals display elevated expression of the fusion control construct but not of IRES-containing constructs. All images are taken under the same conditions. **C**. RT-qPCR analysis of the RNA samples extracted from the animals shown in panel B. Each bar represents an average of 3 biological replicates. The values were first normalized with *rpl12* qPCR counts. Then, the readouts from wild-type animals expressing the fusion control constructs were set to 1. A disparity in the concentrations of mCherry and EGFP sequences in the IRES-containing construct is evident. Primers used for qPCR amplified a 205 bp-long product at 5’ end of the mCherry-encoding sequence, and the 80 bp-long product at the 3’ end of the EGFP-encoding sequence (see Table S1). **D**. The Northern blot was performed using the samples from panels **B** and **C**. RNA resolved in denaturing agarose gel was probed with [α-^32^P]dCTP-labeled fragments corresponding to the mCherry or EGFP CDSs. IRES-containing samples display truncated RNA isoform that accumulates at high levels. The levels of the full-length CrPV IRES-containing transcript were low and did not respond to NMD activity, while the full-length DCV transcript also accumulated to higher levels even in the presence of functional NMD. One representative image of three replicates is shown. FL denotes the full-length transcript, TR denotes the truncated transcript. rRNA was used as a size marker. **E**. Quantification of band intensities from panel **D**. Each bar represents the average of three biological replicates. Samples from wild-type animals expressing the fusion control constructs are set as ones. **F.-G**. RNAi-mediated knockdown of NMD components only slightly affects the expression of IRES-containing constructs. **F**. A crossing scheme of eye-specific RNAi-mediated knockdown of *Upf1* and *Upf2*, the two critical components of the NMD machinery. Wild-type flies were used as a control. **G**. RT-qPCR analysis performed essentially as in panel C with normalization made with Gal4 readouts instead of *rpl12*. Levels of the fusion control construct in the wild-type background were set as one. mRNAs produced from the IRES-containing constructs display only a minor upregulation compared to control. An average of three independent replicates is shown. The minor differences in transcript sensitivity to NMD here and in the panel C might be due to variability in NMD activity between tissues. * - Student’s t-test p-val<0.05, ** - p<0.01, *** - p<0.001. Error bars represent SD. Colors of asterisks corresponds to the experimental group compared: green for GFP, red – mCherry, dark red in parenthesis – truncated isoform of the transcript.

### IRESs induce premature polyadenylation

To determine the potential mechanism of transcript truncation, we tested if shorter transcripts (that comprise most of the mCherry-encoded RNA fraction) are polyadenylated. We purified the poly(A)+ RNAs and analyzed these samples with RT-qPCR. Purification resulted in a ∼200-fold decrease in the amounts of 18S ribosomal RNA (blue bars). However, no significant changes were detected in stoichiometric proportions between mCherry and EGFP-encoding RNAs (**Fig. 3A**). Thus, the truncated mCherry-encoding transcripts are polyadenylated, suggesting the presence of cryptic polyadenylation sites in IRES sequences. To map these potential sites, we performed 3’ - RACE sequencing. This analysis suggested potential polyadenylation signals within both IRESes ∼20 bp upstream of the polyA tail (**Fig. 3B**, triangles). The sequence located in the CrPV IRES perfectly matches the polyA signal consensus AAUAAA (Retelska et al., 2006). Sites found in the DCV IRES share only a partial similarity likely explaining increased amounts of the full-length transcript in DCV samples when compared to CrPV (**Fig. 2C and D**). While a potential polyA site in the CrPV IRES sequence was localized not far from its 5’ end, in the DCV IRES it was found within the 3’ end of the DCV IRES, immediately upstream of the translation start site. This might explain slight differences in mobility of the truncated transcripts shown in **Fig. 2D**, as well as the identification of truncated DCV IRES-containing transcript with EGFP probe.

**Fig. 3.**
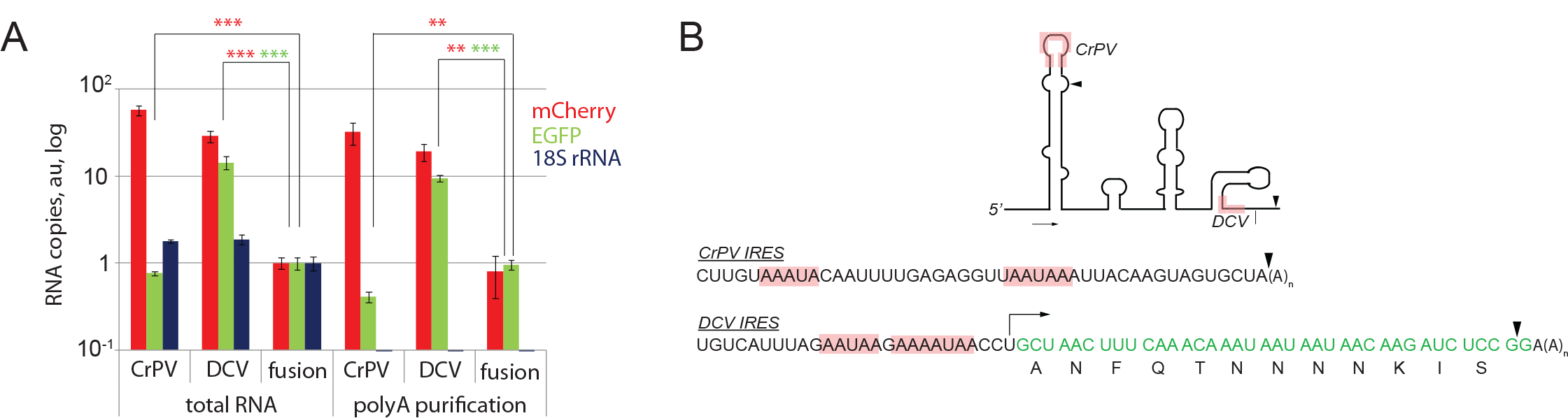
Evidence for cryptic polyA sites in IRES sequences. **A**. Truncated transcripts are polyadenylated as the ratio remains the same upon purification with polyT magnetic beads. Normalization was done as in panel **2C**. An average of three technical replicates is shown. * - Student’s t-test p-val<0.05, ** - p<0.01, *** - p<0.001. Error bars represent SD. **B**. 3’ UTRs were amplified with 3’ RACE and sequenced. Motif analysis suggest the presence of cryptic polyadenylation sites in CrPV and DCV IRESs. The position of potential cryptic polyA sites (consensus sequence AAUAAA) within IRES structures are labeled in pink in the secondary structure image and in the sequence. The triangles mark the start of the polyA stretches in the transcripts found upon sequencing (∼20 bp downstream of the signals). IRES sketches of the figures are from (Jan, 2006) with no pseudoknots shown.

Our model suggests that IRES sequences placed in the intergenic spacer introduce cryptic polyadenylation sites in DNA, leading to the production of truncated transcripts with short 3’ UTRs and resistance to NMD (**Fig. 2, 3**). In this way, the constructs retain the enhancer activity located in the SV40 sequences, but NMD-dependent RNA destabilization is mitigated. Furthermore, potential NMD suppressors located in CrPV IRES sequence could contribute to full-length transcript stabilization (**Fig. 4**).

**Fig. 4.**
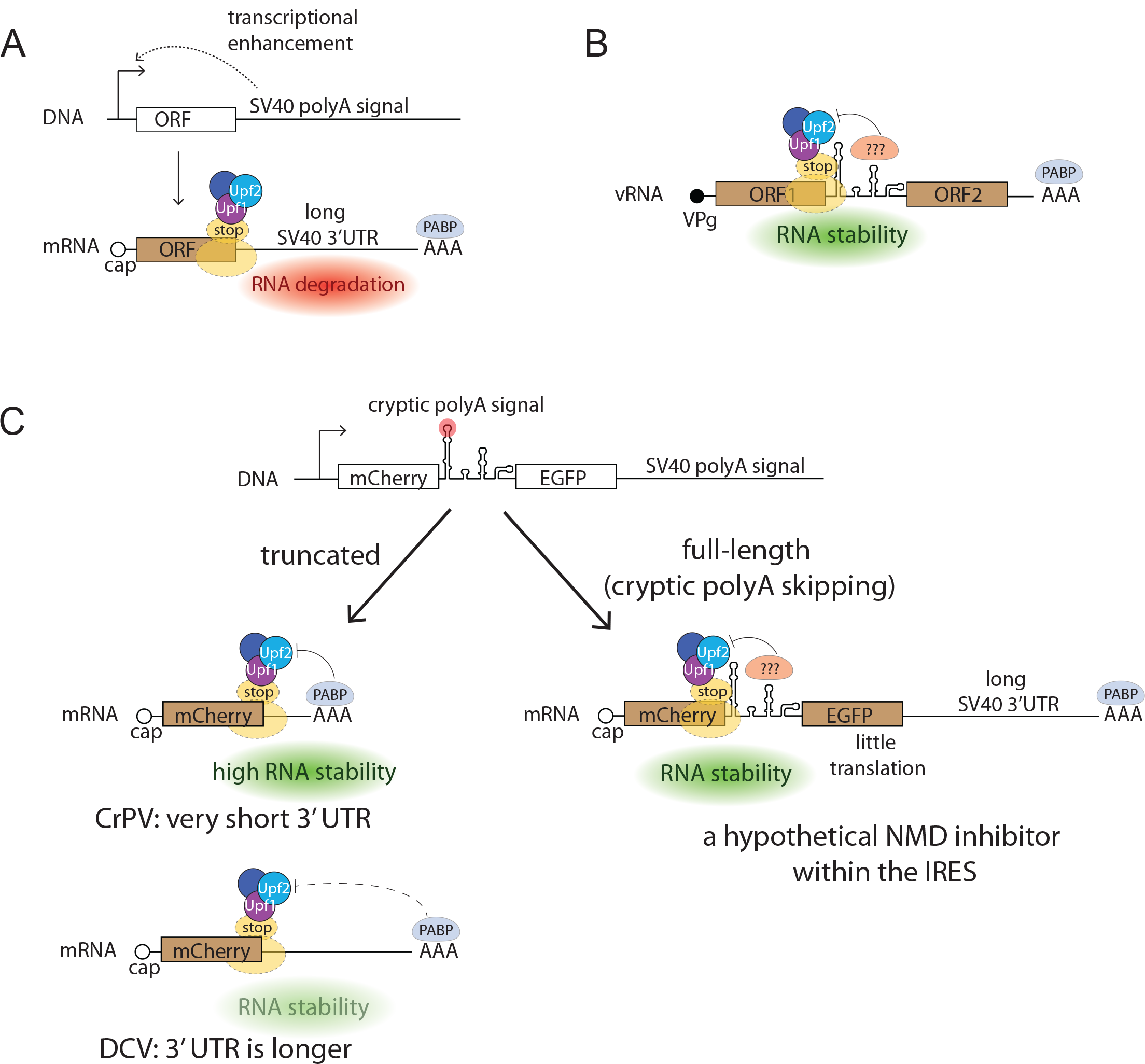
A conceptual scheme of IRES sequence-mediated overexpression. **A**. SV40 polyadenylation signal enhances transcription but also makes the transcript sensitive to NMD. **B**. Intergenic IRES position within the virus genome provides an evolutionary rationale for the presence of NMD inhibitors within the IRESs. **C**. Cryptic polyadenylation signals within the IRESs result in transcript truncation and stabilization that is dependent on 3’ UTR length. Putative NMD suppressor sequences might reside within IRESs, thus explaining decreased sensitivity of the full-length constructs to NMD.

## Discussion

The experiments described above suggest that CrPV and DCV IRES sequences present in bicistronic constructs induce mRNA stabilization and a robust increase in production of the protein encoded by the 5’ proximal cistron. Our data have led us to suggest the following model (**Fig. 4**). In the absence of IRESs, the 3’ end SV40 sequence plays a dual role in gene expression: it enhances transgene transcription and destabilizes mRNA due to increased sensitivity to NMD (**Fig. 4A**).

Earlier, Metzstein and Krasnow found that transgenic proteins expressed using the cassettes with the SV40 polyadenylation element were more abundant compared to those expressed with Hsp70 polyA site. They also observed that transgenes derived from SV40 polyA but not Hsp70 polyA signal-containing constructs were upregulated in NMD-null “photoshop” mutant flies (Metzstein and Krasnow, 2006). A potential mechanism for mRNA destabilization, mediated by the SV40 sequence, is related to its elongated 3’ UTR.

In mammals, premature stop codons are detected by translation termination upstream of an exon-exon junction, leading to NMD-dependent degradation (Ivanov et al., 2008). An alternative mechanism relies on the 3’ UTR length. In mammals it operates in addition to exon-exon junction detection, while in insects, where many genes lack introns, 3’UTR length is the main mechanism for premature stop codon detection by NMD (Behm-Ansmant and Izaurralde, 2006; Nogueira et al., 2021). It was shown that activation of NMD machinery by terminating ribosomes is blocked by polyA-binding protein (PABP) (Ivanov et al., 2008). It is believed that lengthy 3’ UTRs bring PABP away from the stop codon, removing this block and destabilizing the transcript. Consistently, truncating the SV40 derived sequence, but not removing the intron located within it, affects sensitivity to NMD in UAS/SV40 constructs (Nelson et al., 2018).

In addition, the full-length bicistronic transcripts containing IRESs showed decreased sensitivity to NMD compared to the control fusion construct. This effect was especially pronounced in CrPV IRES encoding constructs. The corresponding EGFP coding sequence (CDS)-specific RT-qPCR readouts showed substantially lower elevation (**Fig. 2G**) in NMD-deficient strains, unlike those in samples with the fusion control construct. Northern blot and RT-qPCR analyses showed that the full-length transcript derived from the CrPV IRES-containing cassette was negligibly affected by *Upf2* inactivation (**Fig. 2C, D-E, G**). This result suggests that the CrPV IRES-containing bicistronic transcripts have an increased resistance to NMD if compared to control. Based on these results we hypothesize the existence of NMD inhibitor within CrPV IRES sequence as the most parsimonious explanation of the observed effect. The DCV IRES containing construct showed elevated levels of mCherry and EGFP RNA in the presence of active NMD. At the same time, these levels increased when NMD was inhibited (**Fig. 2C, D-E, G**). We hypothesize that this effect results from a combination of a weaker polyadenylation site with a weaker NMD suppressor present in the IRES. Further research is needed to tests these hypotheses and elucidate the mechanisms of interactions between IRESs and NMD.

In dicistrovirus genomic RNAs, intergenic IRESs are located immediately downstream of the first CDS which encodes non-structural proteins (Bonning and Miller, 2010). Termination of translation at the stop codon of this CDS should activate NMD machinery due to its long distance from the polyA (Behm-Ansmant and Izaurralde, 2006). Growing evidence suggests that NMD is also involved in the degradation of many RNA(+) virus genomes (Balistreri et al., 2014; Contu et al., 2021; Fontaine et al., 2018; Ramage et al., 2015; Withers and Beemon, 2011). The NMD-inhibiting cis-acting RNA structures which are located immediately after stop codons were previously described in retroviruses (Balagopal and Beemon, 2017; Barker and Beemon, 1994). The intergenic IRES sequences that take positions downstream from the ORF1 stop codons might possess an additional activity to suppress NMD (**Fig. 4B**). Further research is required to validate these suppressors and find the mechanisms of their action.

We suggest that dicistrovirus IRES sequences, when placed into artificial bicistronic constructs, may stabilize mRNA transcripts by (i) shortening the 3’ UTR through cryptic polyadenylation signals at the DNA level and (ii) directly inhibiting NMD machinery with virus-specific suppressing elements at the RNA level.

It should be noted that neither CrPV nor DCV IRESs mediate any detectable translation of EGFP, suggesting that they are not efficient translation activators (**Fig. 1B**). This aligns with the observation that intergenic IRES activity is low at early stages of CrPV infection, but increases in the middle of the replication cycle due to direct or indirect stimulation by viral ORF1-derived protein products (Khong et al., 2016). Experiments in flies corroborate the model suggesting that host’s translation shut-off might serve as a mechanistic switch activating ORF2 translation (Lidsky et al., 2023).

In summary, the combination of the transcription-enhancing SV40 polyadenylation signal and the IRESs, which introduce cryptic polyadenylation sites and result in truncation of the 3’ UTR may constitute a novel, effective approach for high-level protein production in *Drosophila*. Expression of mCherry at high levels with the use of this system was compatible with normal fly development (**Fig 1B**) and showed no apparent eye phenotypes (**Supplementary Fig 1C**) indicating low toxicity of the constructs. Previous methods for tuning expression levels in flies were based on multimeric transcription factor binding sites (Pfeiffer et al., 2010), translational regulators (Pfeiffer et al., 2012; Zufferey et al., 1999), or enhanced mRNA nuclear export (Huang and Gorman, 1990). Our approach is unique in its focus on increased mRNA stability. It can be applied not just to fluorescent markers, but potentially to other reporters and biotechnological protein production.

## Materials and methods

### Cloning and injection

*pUASt-attB* vector, which contains a 694 bp-long fragment with SV40 polyadenylation signal (Bischof et al., 2007) was used for cloning to create IRES-encoding constructs. To obtain *pUASt-mCherry-CrPV_ires-EGFP-attB* and *pUASt-mCherry-DCV_ires-EGFP-attB* plasmids, encoding the IRES-containing bicistronic RNAs, *pUASt-attB* vector was digested with EcoRI and XbaI enzymes. Fragments encoding EGFP and mCherry were amplified by PCR. EGFP fragment was digested with BglII and XbaI. mCherry fragment was treated with EcoRI and BsrGI. IRES sequences were synthesized by Genescript and excised from their vector plasmid with BsrGI and BglII. The constructs were then injected into M{3xP3-RFP.attP}ZH-86Fb docking site strain (Bischof et al., 2007) by BestGene Inc.

### Fly lines

The *Upf2*^*25G*^/FM7 flies (Metzstein and Krasnow, 2006) were a kind gift of Prof. Mark M. Metzstein (University of Utah). A ubiquitous *daughterless*-*Gal4* driver, *GAL4-da*.*G32* was on the third chromosome (Wodarz et al., 1995); in the text, it is referenced as *da-Gal4. eyeless-Gal4* expression system, *y, w, ey-Flp, Act5c>CD2>Gal4* chromosome (Pignoni and Zipursky, 1997) was ascribed in the text as *ey>Gal4* and was kindly provided by Prof. Hugo Stocker (ETH, Zurich). Gal4 drivers did not contain SV40-polyA sequences. *y, w* flies (kindly donated by Prof. Stefan Lusching, University of Muenster) were used as a wild type control. *TRiP*.*GL01485* and *TriP*.*JF01560* lines used to knockdown *Upf1* and *Upf2* correspondingly were received from the Bloomington stock center. GAL4 under control of the Hsp70 promoter and two copies of the sevenless enhancer (Ruberte et al., 1995) was used to validate RNAi constructs. UAS-GCN2^Act^ (Bjordal et al., 2014) insertion was generously provided by Prof. Pierre Leopold (Institut Curie). Fly stocks were kept at 18°C, while the crosses – at 25°C.

### Sample preparation, microscopy, and image analysis

Whole flies, larvae, and fly eyes were imaged with Nikon SMZ1500 and Leica MZ16 F microscopes and deconvolved with Helicon Focus software (HeliconSoft).

3^rd^ instar larvae eye inmaginal discs were dissected in PBS, fixed with 4% paraformaldehyde on PBS for 20 minutes at room temperature, washed with PBS, quenched with 1M glycine on PBS, washed again, and mounted with Vectashield media (Vector Laboratories). Confocal images were acquired with Nikon Ti Spinning Disc confocal microscope equipped with 20x objective and processed using Imaris software (Bitplane).

### RNA isolation and RT-qPCR

Total RNA isolation was performed with TRI-reagent (Sigma) as per the manufacturer’s recommendations. Upon isolation, samples were treated with DnaseI (New England Biolabs). The polyA-enrichment was performed with NEXTflex™ Poly(A) kit purchased from BIOO Scientific. RT reactions were performed with Maxima RT (ThermoFisher Scientific), while qPCR reactions were performed with SensiFAST™ SYBR® No-ROX Kit (Bioline). For oligonucleotide sequences, see Table S1. Ten animals were used per replica. Three biological replicates were used for the experiments shown in panels **2C, D, E, F**. Panel **3A** is a representative result of three independent repeats. **3’UTR sequencing** was performed with 2 round of PCR after RT reaction carried out with the use of polyT oligonucleotide. In the first round, the polyT-containing reverse oligo with 5’ adaptor sequence and a forward oligo annealing within mCherry sequence were used. The PCR product was excised from the agarose gel and used as a template for the second round. A nested pair consisted of a reverse oligo annealing to the adapter sequence and a forward oligo annealing in the mCherry region. The resulting products were used for Sanger sequencing.

**Northern blot** was performed as described (Saleh et al., 2009). Electrophoresis was performed on 10 μg of total RNA per sample in 1% (w/v) agarose gels containing 1.1 mM formaldehyde. The RNA was transferred overnight by capillarity to an Amersham Hybond-N+ membrane (GE Healthcare Life Sciences) and covalently bound to the membrane using a Stratalinker UV crosslinker. Northern blots were hybridized with DNA probes generated by a random-primed labeling reaction with use of Amersham Ready-To-Go DNA Labelling Beads (GE Healthcare Life Sciences) supplemented with [α-^32^P]dCTP and a corresponding PCR fragment as a template. Membranes were exposed overnight to a Phosphor Imager screen at room temperature. Bands were quantified with Fiji software (Schindelin et al., 2012).

## Data availability statement

Fly stocks and plasmids are available upon request. The authors affirm that all data necessary for confirming the conclusions of the article are present within the article, figures, and tables.

## Acknowledgements

S.E.D. is a member of the Interdisciplinary Scientific and Educational School of Moscow University “Molecular Technologies of the Living Systems and Synthetic Biology”. We thank Profs. Mark M. Metzstein, Hugo Stocker, and Stefan Lusching for sharing fly stocks.

## Figure legends

**Supplementary Fig 1. A-B**. Validation of RNAi constructs. Control *yw* flies or flies carrying UAS-RNAi constructs targeting Upf1 (*TRiP*.*GL01485*, **A**) and Upf2 (*TriP*.*JF01560*, **B**) were crossed with the flies encoding Gal4 under control of heat shock and sevenless promoter sequences. This driver was used to produce weaker knockdown of these NMD components that would not interfere with fly viability. Adult males of the resulting crosses were collected for RNA extraction and qPCR. Both RNAi constructs displayed statistically significant downregulation of the corresponding mRNA. Four biological replicates were used for each experiment. **C**. Eyes of the flies analyzed in the **Fig 2F-G**. No strong phenotypes observed.

